# Cell-autonomous BMP signalling plays a key role in the maintenance of tumour cell EMT and migration programs in human ovarian carcinoma

**DOI:** 10.1101/2023.04.30.538847

**Authors:** George Joun, Fatemeh Zolghadr, Priyanka Chakraborty, Thi Yen Loan Le, James J.H. Chong, Australian Ovarian Cancer Study Group, David D. L. Bowtell, Anna DeFazio, Mohit Kumar Jolly, Naisana Seyedasli

## Abstract

Epithelial-mesenchymal transition (EMT) plays a key role in tumour initiation, metastasis and resistance to therapy. Cells undergoing EMT, assume multiple semi-stable transitional states along the epithelial-mesenchymal axis that necessitates tight regulatory cascades. Although more is known about pathways involved in the initial induction of EMT, cascades that mediate/maintain the transitional states and/or the final mesenchymal phenotype are yet to be elucidated. In this study, we have assessed the role of bone morphogenic protein (BMP) signalling pathway in the regulation of cancer cell EMT and migration. Mining existing data from ovarian carcinomas, we defined the BMP pathway among the key pathways enriched in tumours with elevated EMT signatures, with a significant correlation in the expression of EMT markers with BMP ligands and downstream targets of the BMP pathway. Functional inhibition of the BMP pathway in ovarian cancer cells by a small molecule inhibitor, DMH1, resulted in impaired migration and depressed EMT signatures in both *in vitro* and *in vivo* models. Finally, ectopic induction of EMT in ovarian cancer cells through activation of an independent pathway, TNFα, resulted in the selective induction of Smad-mediated BMP pathway suggesting a role in maintenance of EMT, secondary to EMT induction.

## Introduction

Epithelial mesenchymal transition (EMT) is an evolutionarily conserved process actively involved in embryonic development and morphogenesis as well as adult homeostasis and disease [1–5]. In carcinoma, EMT plays a key role in the neoplastic transformation of the epithelial tissue and acts at multiple stages of tumour initiation, progression and metastasis [6, 7]. EMT has as well been described as a key component in the induction and maintenance of tumour initiating or cancer stem cells (CSC) [8, 9] with major roles in tumour resistance to therapy [10]. During EMT, cells within the compact structured epithelium gain properties of mesenchymal cells including motility and minimal dependence on cell-cell contacts. Evidence confirms cellular states along the epithelial-mesenchymal (E/M) axis exist in a continuum with defined E/M signatures [11, 12]. The (semi)-stable states of cells along the E/M axis suggest the presence of tightly regulated upstream signalling networks [13, 14]. Numerous signalling pathways are suggested to act upstream of EMT as an inductive cue [15, 16], however the insight into pathways involved in the maintenance of transitional EMT states is minimal [17]. Components of the bone morphogenic protein (BMP) pathway have been shown to be involved in the induction of EMT in different cancer types [18–20]. In this study, we highlight a key role for the BMP pathway in the maintenance of EMT downstream of its induction.

BMP signalling acts as a master regulator throughout early embryonic development and tissue morphogenesis [21, 22] with continued regulatory roles in adult tissues [23–25]. BMPs are members of the transforming growth factor-β (TGF-β) family, which bind several target receptors including BMP receptors 1A, 1B and 2 [23, 26]. Upon ligand-receptor binding, receptor multimerisation and downstream phosphorylation events lead to transcriptional regulation of target genes [23, 26]. BMP pathway activity is commonly mediated through phosphorylation of a core complex of proteins known as small mothers against decapentaplegics (SMAD) (Smad1/5/9) that upon binding to the co-Smad molecule Smad4, can translocate to the nucleus and act on target promoters for transcriptional regulation. Evidence as well support non-canonical molecular mediators downstream of activated BMP receptors including phosphatidylinositol 3-kinase (PI3K)/AKT, mitogen-activated protein kinase (MAPK), nuclear factor kappa B (NF-κB) and Janus kinase/signal transducer and activator of transcription (JAK/STAT) pathways [23, 26–29]. BMP pathway regulates tumour cell properties including cell proliferation [22, 30, 31], invasiveness and metastasis [32]. Although multiple studies highlight a role for the BMP signalling pathway in the induction of EMT [33, 34], less is known about its involvement in the transitional states, or the maintenance of the mesenchymal phenotype. In this study, we highlight a key role for the BMP signalling pathway downstream of EMT induction, as a cell-autonomous pathway involved in the maintenance of the mesenchymal state.

## Materials and Methods

### Cell lines, general tissue culture, stable transfection of cell lines

AOCS15 and COLO316 [35] are high-grade serous ovarian cancer (HGSC) cell lines. AOCS15 is a new, patient-derived cell line from the Australian Ovarian Cancer Study (AOCS) and was established in Prof David Bowtell’s laboratory from malignant ascites obtained from a 62-year-old woman diagnosed with FIGO stage IV, high-grade serous ovarian carcinoma (manuscript in preparation). Both cell lines were authenticated by STR profiling and were routinely tested and shown to be free of *Mycoplasma* contamination. Cells were cultured in standard culture medium (RPMI1640, Sigma) containing 10% fetal bovine serum (Gibco), 20 mM HEPES (Gibco) and 2 mM Glutamax (Gibco). The cells were incubated at 37°C in 5% CO2.

To generate AOCS15 cell lines with stable expression of GFP and mCherry, HEK293T cells were transduced with pUE-Lenti-eGFP or pUE-Lenti-mCherry plasmids and the lentiviral packaging plasmids pMD2 VSV-G, pMDLgp RRE and pRSV REV using calcium phosphate precipitation as described before [36]. After 72 hours of transfection, the viral supernatants were collected and centrifuged at 3000 rpm for 1-2 hours at 4°C, and then used to transduce AOCS15 cells. The cells were transduced with lentivirus in media containing polybrene (8 µg/ml) for 24 hours followed by antibiotic selection with 10 µg/ml Blasticidin (Thermo Fisher Scientific, Cat# A1113903). Non-transduced cells were included as controls. Successful eGFP and mCherry over-expression was confirmed by fluorescence microscopy.

### Protein isolation, Western Blot analysis, immune-cyto/histochemistry, treatment with BMP pathway inhibitor and blocking with monoclonal antibodies

Total protein from COLO316/AOCS15 was isolated and transferred to nitrocellulose membrane using standard protocols [37]. Presence of active BMP signalling pathway was detected by an antibody against the phosphorylated form of Smad1/5/9 protein complex (Cell Signalling Technology, Cat#13802S) or pSmad1/5 (Thermo Fisher Scientific, Cat#700047) as indicated in figures. Total Smad1 was detected with an antibody against this protein (Cell Signalling Technology, Cat# 9743). Total Histone2B (H2B; Abcam, Cat #ab18977) or Vinculin (Sigma Aldrich, Cat#V9131) were used as loading controls. To demonstrate the expression of pSmad1/5/9, the protein was detected in PFA-fixed COLO316/AOCS15 cells using standard immune-cytochemistry protocols [37]. To detect mCherry and e-GFP from transplanted AOCS15 cells in the CAM migration assay, membranes were fixed and processed for immunohistochemistry using standard protocols. Cells were detected using antibodies against mCherry (Abcam, Cat# ab183628) and GFP (Abcam, Cat# ab5450). All imaging was performed on Olympus BX60 fluorescent microscope. To inhibit the BMP pathway activity, DMH1 (Sigma-Aldrich, Cat# D8946) was reconstituted in DMSO and added to the culture medium in the indicated concentrations. To block BMP6, 5 ug of total BMP6 monoclonal antibody (Thermo Fisher Scientific, Cat# MA5-23844) was added to the culture medium for 48hrs and cells were then harvested for protein analysis. Untreated and IgG-treated cells were used as controls. For induction of EMT, cells were treated with 10 ng/ml of the recombinant human TNFα protein (Merck-Millipore, Cat# GF023) and the outcome was assessed at RNA or protein levels.

### TCGA data analysis, calculation of EMT scores, statistical analysis of expression correlates

The TCGA ovarian cancer microarray dataset was downloaded from UCSC xena tools and HGSC samples were selected based on the *TP53*, *NF1*, *BRCA1*, *BRCA2*, *RB1* and *CDK12* mutation status [38]. EMT scores were calculated using two methods, KS-statistic (two sample Kolmogrov-Smirnov test) [39] and MLR (multinomial logistic regression, George, 2017 #6501}. The KS-statistic score varies on a scale of -1 to 1, whereas the MLR method-based model quantifies the EMT status of a given sample on a scale of 0 to 2. In both scoring methods, samples with the higher scores correspond to more mesenchymal phenotype. All samples were segregated into high and low group based on the EMT score threshold of 0. To verify the variation between EMT high and low groups differential gene expression (DEG) analysis was performed using limma bioconductor R-package. Next, DEGs were used as an input to DAVID database for pathway and gene ontology analysis, and p<0.05 and gene count >2 were set as the cut-off point. A total of 2695 DEGs, which were significantly enriched in biological processes, were screened.

### Real-time RT-PCR, Semi-quantitative RT-PCR, Statistical analysis

Total RNA was extracted from COLO316 cells using Bioline Isolate II RNA Mini Kit (BIO- 52073) and cDNA synthesis was carried out using Bioline Sensifast cDNA synthesis kit (BIO- 65054), following manufacturer’s instructions. Real-time PCR was carried out using Bioline SensiFAST SYBR Lo-ROX Kit (BIO-94005) and reactions were run on Agilent Technologies Stratagene Mx3000P machine with data analysed on Agilent MxPro qPCR software. The primer sequences were as outlined in Table 1. Data analysis was performed using the standard ι1Ct/Cp method. Final fold change was calculated relative to normalised ι1Ct levels in human induced pluripotent stem cell cDNA as control for an epithelial tissue or to the control vehicle-treated samples in case of TNFα−treated cells.

For semi-quantitative RT-PCR, total RNA and cDNA were synthesised as described above. PCR amplification was performed using BIOLINE MyTaq PCR kit (cat no. BIO-21111) with specific primers designed for BMP genes as listed in Table1. Human universal total RNA (Agilent) was used as a positive control. Amplification products were visualised by electrophoresis on 1.5% w/v agarose gel and visualised by Gel Red (Biotium, Cat. No. 41002) using manufacturer’s instructions.

All statistical significance was determined using the standard Student’s t-test (Microsoft Excel), or one-way ANOVA (GraphPad, Prism) based on the average signal between biological triplicates. In all cases, p-value<0.05 was regarded as significant.

### In vitro cell migration and in ovo chorioallantoic membrane (CAM) migration assays

For *in vitro* cell migration assays, cells were seeded in co-culture stamping devices as described before [40]. Four hours post seeding, the stamps were removed and cell strips were covered with control/vehicle-treated or DMH1-containing culture medium. Cells were then imaged at 15 minute intervals on the Leica Live Cell Imaging (DMI6000B) microscope for a total period of 24 hours. Four corresponding time points between control and DMH1-treated cells were selected for the analysis of cell migration. At each time point, the cell-free area (CFA) between cell strips was measured by ImageJ [41] and normalised to CFA at time 0. Relative CFA was then compared at given time points between control and DMH1-treated cells.

For the *in ovo* CAM migration assay, AOCS15 cells with stable expression of mCherry (AOCS15^mCherry^) and eGFP (AOCS15^eGFP^) were transplanted to the CAM of E3 (3 days post-incubation) chicken embryos. To block the endogenous BMP signalling activity, AOCS15^eGFP^ cells were treated with 500 nM DMH1 for 48hrs prior to transplantation while vehicle-treated cells were used as control. For transplantation, cells were harvested and resuspended in culture medium as 6000 cells/ul. AOCS15^mCherry^ and AOCS15^eGFP^ (Control/DMH1-treated) cells were then mixed in 1:1 ratio. Cell mixture was further mixed 1:1 with Matrigel (Corning, Cat#354230) and formed into 30 ul hanging drops with further incubation for 30 min for matrigel to solidify. Meanwhile, the CAM from E3 chicken embryos was exposed through introduction of a small opening in the eggshell. Matrigel drops containing AOCS15^mCherry^ + AOCS15^eGFP-control^ and AOCS15^mCherry^ + AOCS15^eGFP-DMH1^ were then placed on the CAM in individual eggs and immobilised by covering with a round 13 mm coverslip. Eggs were then closed and re-incubated in humidified incubators at 37°C for an additional 72 hrs. The membranes were then dissected in cold PBS, fixed in 4%PFA/PBS for 30 min at room temperature and processed for standard cryo-embedding and immuno-histochemistry.

## Results

### Scoring of ovarian tumour samples based on levels of EMT, highlights multiple signalling pathways enriched in EMT^high^ tumours

Several signalling pathways have been associated with the EMT program in ovarian cancer [42]. The majority of these studies however, have focused on the role of individual signalling molecules/pathways in the induction of EMT [43], with limited insight into the pathways upstream of EMT maintenance. To determine whether signalling pathways are differentially associated with levels of EMT, we used an EMT scoring algorithm (Kolmogorov-Smirnov test- KS) [39] to stratify 346 high grade serous ovarian carcinoma (HGSC) tumour samples from the cancer genome atlas (TCGA) into EMT^high^ and EMT^low^ based on the EMT score threshold of 0. The EMT score varies on a scale of -1 (fully epithelial) to + 1 (fully mesenchymal) (Figure S1A). We first verified that the levels of EMT transcription factors *ZEB1*, *ZEB*2, *SNAI2*, *TWIST1* and the canonical mesenchymal marker, Vimentin (*VIM*) [44], were higher in EMT^high^ group (Figure S1B). In contrast, the levels of mesenchymal epithelial transition (MET)-inducing transcription factors *GRHL2* [45] and *OVOL2* [46] and the epithelial splicing regulators *ESRP1/2* [47] were higher in EMT^low^ group (Figure S1B). The same samples were then re-analysed by EMT scores calculated via another EMT scoring method based on multinomial logistic regression (MLR) [48]. A positive significant correlation was observed between KS and MLR scores (Figure S1C). As anticipated, a similar pattern of distribution was observed in samples segregated by MLR scores, with mesenchymal markers enriched in EMT^high^ and epithelial markers enriched in EMT^low^ samples (Figure S1D) while constant negative correlations were observed between various pairwise epithelial and mesenchymal markers in EMT^high^ and EMT^low^ (Table S2). To highlight signatures of EMT^high^ and EMT^low^ groups, differentially expressed genes (DEGs) were analysed. Clustering of these samples based on 100 most differentially expressed genes resulted in 3 clusters, Cluster 1 – EMT^high^, Cluster 2- EMT^low^ and Cluster 3 – mixed group (Figure 1A). Next, DEGs were used as an input to the DAVID database for pathway and gene ontology analysis (Figure 1B). Through the gene ontology function, significantly enriched pathways were identified, among which signalling pathways mediated by Semaphorin-plexin, Wnt, BMP, TGF-β and Integrin were significantly enriched in EMT^high^ vs EMT^low^ (Figure 1B). Enrichment of these pathways in tumours with elevated EMT signatures suggests a role for these signalling pathways in the regulation of the EMT network.

**Figure 1.**
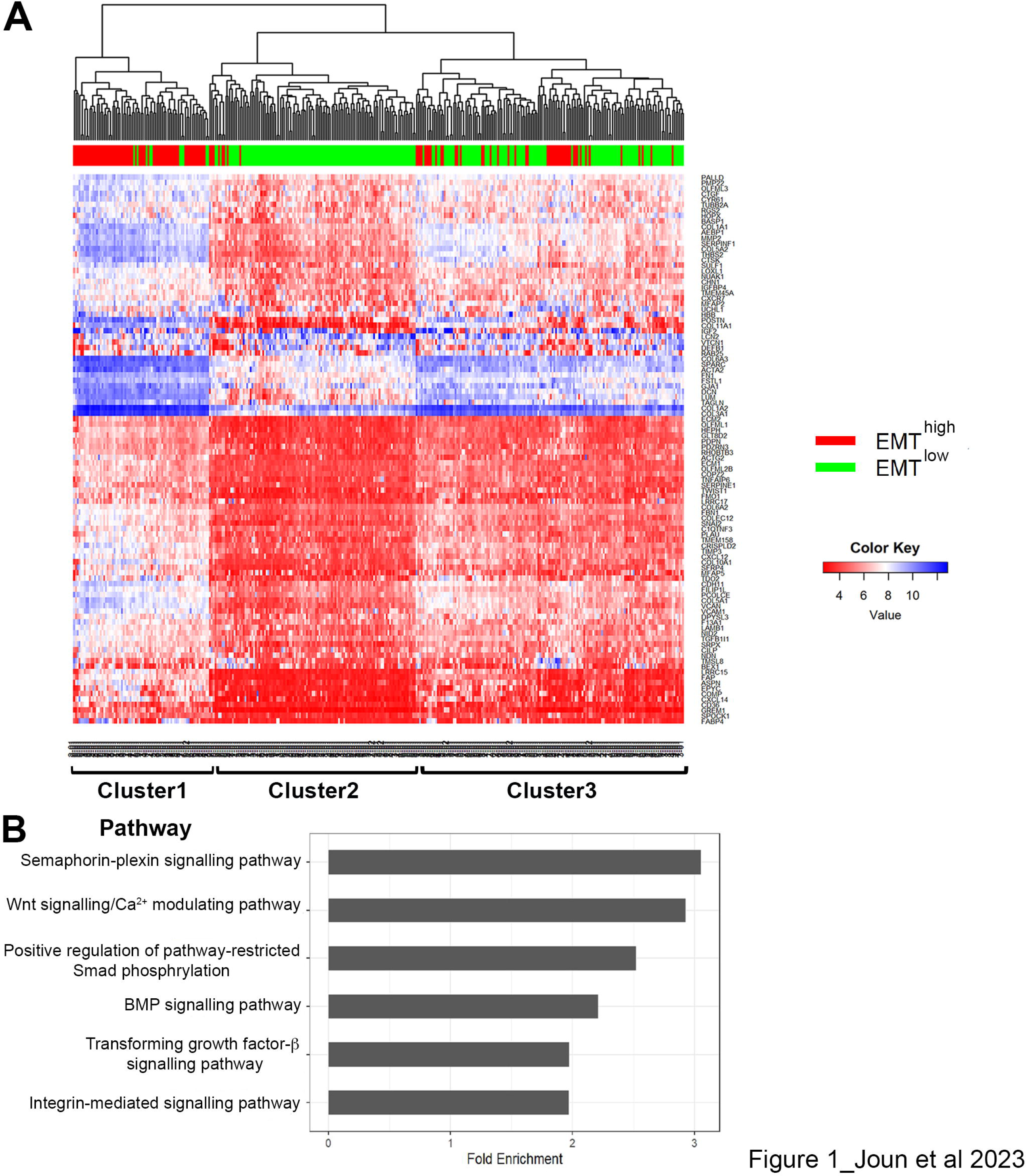
Enrichment of signalling pathways in ovarian tumours with elevated EMT signatures. A) Clustering and heat-map for the expression of top 100 differentially expressed genes from TCGA HGS tumours with low (EMT^low^) and high (EMT^high^) EMT scores [39]. B) Signalling pathways enriched among differentially expressed genes in EMT^high^ vs EMT^low^ tumours.

### BMP pathway ligands and downstream target genes are enriched in tumours with elevated EMT signatures

Following the pathway analysis (Figure 1B), we investigated the association between the expression levels of immediate genes involved in the BMP signalling pathway activity and the expression of key EMT markers [49] in TCGA HGSC data. Correlations between the expression of BMP ligands (BMP1-15), receptors (BMPR1A, 1B and 2), and the global downstream BMP target genes, the inhibitors of differentiation (ID) [50], with EMT transcription factors in TCGA HGS data were analysed. The expression levels of two BMP ligands, *BMP1* and *BMP4*, and three ID genes, *ID1*, *2* and *3* with multiple EMT transcription factors including *ZEB1*, *ZEB2*, *SNAI2* (Slug) and *TWIST*1, and the mesenchymal marker, Vimentin (Figure S2) were positively associated. Further, scoring of HGSC samples based on EMT signatures using either KS or MLR methods, showed significantly enriched expression of *BMP1* and *BMP4* in EMT^high^ tumours (Figure 2A and B). Likewise, enriched expression of *ID1*, *ID2* and *ID3* was observed in EMT^high^ tumour subset using the KS method, although the MLR-based scoring method only showed a significantly-enriched expression for *ID3* in EMT^high^ tumour samples (Figure 2C and D). Altogether, these findings strongly suggest that BMP signalling pathway plays a key regulatory role in the EMT network with components associated with enhanced EMT signatures.

**Figure 2.**
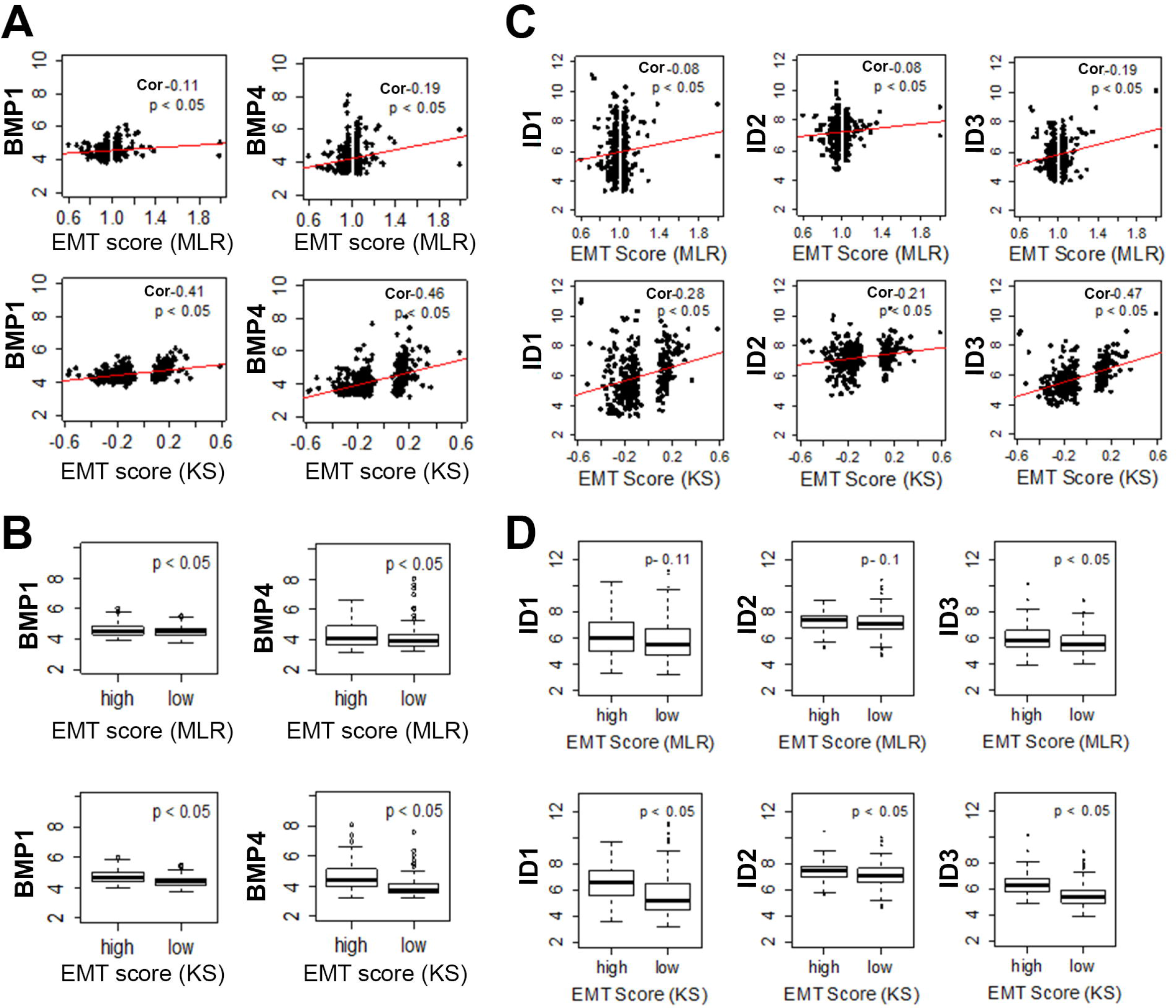
Correlation between BMP pathway components and EMT markers in TCGA data from HGS ovarian tumours. A) The relationship between the expression profiles of BMP1 and BMP4 to EMT scores defined by a linear regression line (red), correlation coefficient (R) and p-value (p) reported at the top-right corner in each plot. Scores were calculated using two scoring methods (MLR vs KS). B) BMP1 and BMP4 expression in subset of samples with high and low EMT scores. (C) The relationship between the expression profiles of ID1, ID2 and ID3 to EMT scores defined by a linear regression line (red), correlation coefficient (R) and p-value(p) reported at the top-right corner in each plot. (D) ID1, ID2 and ID3 expression in samples with high and low EMT scores. p-values based on calculation via students’ t-test.

### Inhibition of BMP pathway negatively affects tumour cell EMT and migration programs

BMP pathway has been shown to act upstream of cancer cell EMT in different cancer types [18–20]. Other studies have however shown a dual, context-dependent role for the pathway with evidence to support both EMT-inductive and repressive roles [24, 25]. To confirm the function of the BMP pathway activity in our HGSC cell lines COLO316 and AOCS15, we specifically inhibited the pathway using a small molecule inhibitor, DMH1 [51, 52]. Treatment with increasing concentrations of DMH1, resulted in significant repression of the BMP pathway activity in the cells as marked by the reduction in phosphorylated Smad 159 (pSmad 1/5/9) protein levels (Figure S3A). We then measured the expression of EMT transcription factors *SNAI1*, *SNAI2*, *TWIST1* and *ZEB1* in response to increasing concentrations of DMH1. Both cell lines had high baseline EMT signatures, marked by comparable levels of *SNAI2*, *TWIST1* and *ZEB1* transcripts to human dermal fibroblasts (HDF) (Figure S3B). Expression of all EMT transcription factors significantly declined in response to higher (500 nM) concentrations of DMH1, while *SNAI2* and *ZEB1* in COLO316 and *SNAI1* in AOCS15 were also significantly repressed upon treatment with the lower (200 nM) DMH1 concentration (Figure 3A). Neither of the treatment regimens, however, could retrieve the expression of the epithelial marker E-cadherin (*E-CAD*) suggesting the absence of a complete MET.

**Figure 3.**
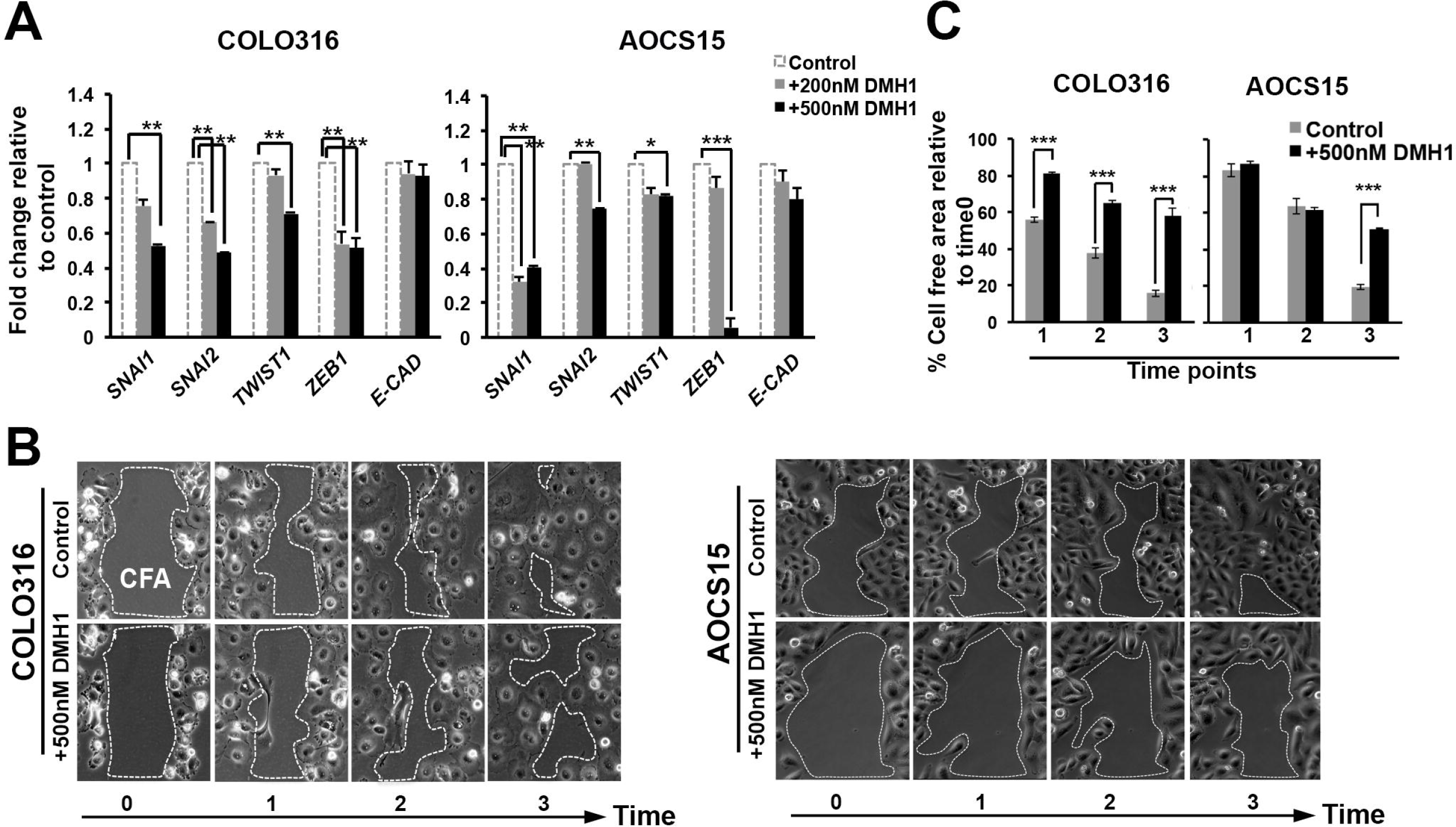
Negative impact of BMP pathway inhibition on EMT and tumour cell migration. A) Real-time RT-PCR analysis of EMT markers in total RNA from two ovarian carcinoma cell lines treated for 48 hrs with increasing concentrations of DMH1, compared to control/vehicle- treated. B) Migration assay comparing the cell free area (CFA) of control/vehicle vs DMH1- treated cells at multiple corresponding time points within 24hrs. C) Quantification of images in (B) (E-CAD: E-cadherin).

Given the significant negative impact of BMP pathway inhibition on the expression of EMT transcription factors, we asked if the effect would functionally translate to impaired tumour cell migration. For this, COLO316 and AOCS15 cells were seeded in specialised cell migration devices [40], treated with control/vehicle or 500 nM DMH1-containing medium and imaged for 24 hours (Figure 3B and C). Corresponding time points between control and DMH1 treatment were then selected for analysis. The analysis of relative cell free area (CFA) between control and DMH1-treated cells showed a significant increase in DMH1-treated CFA at all analysed time points in both cell lines demonstrating a marked delay in cell migration and gap closure, once the BMP pathway was inhibited. Altogether, these results confirm a role for the BMP pathway in positive regulation of the EMT and migration programs in both tested ovarian cancer cell lines.

### A novel in ovo competitive migration assay confirms impaired migration capacity in ovarian carcinoma cells upon BMP pathway inhibition

To address the effect of BMP pathway inhibition in a physiologically-relevant microenvironment, we developed a new migration assay using the chorionic epithelium of the chorioallantoic membrane (CAM) in developing chicken embryos (Figure S3C). AOCS15 cells were genetically tagged with mCherry (red) or eGFP (green) fluorescent proteins. For the migration assay, differentially labelled cells were harvested and mixed in a 1:1 ratio into Matrigel hanging drops (Figure 4A). Matrigel drops containing the control or treated cells were implanted on the CAM of developing chicken embryos (E2) and the eggs were further incubated for 72 hrs before the analysis. Both cell types (AOCS15^mCherry^ and AOCS15^eGFP^) were equally able to migrate along the epithelial layer in the CAM during the 72 hrs post-implantation (Figure S3D). Treatment of AOCS15^eGFP^ cells with DMH1 for 48 hrs prior to implantation (AOCS15^eGFP-DMH1)^ however, resulted in a significant decline in the migration capacity of these cells compared to both the co-transplanted AOCS15^mCherry^ or the control vehicle-treated AOCS15^eGFP-CONTROL^ cells (Figure 4B). This observation further confirmed a key role for the active BMP signalling pathway in the maintenance of migration capacity in ovarian cancer cells.

**Figure 4.**
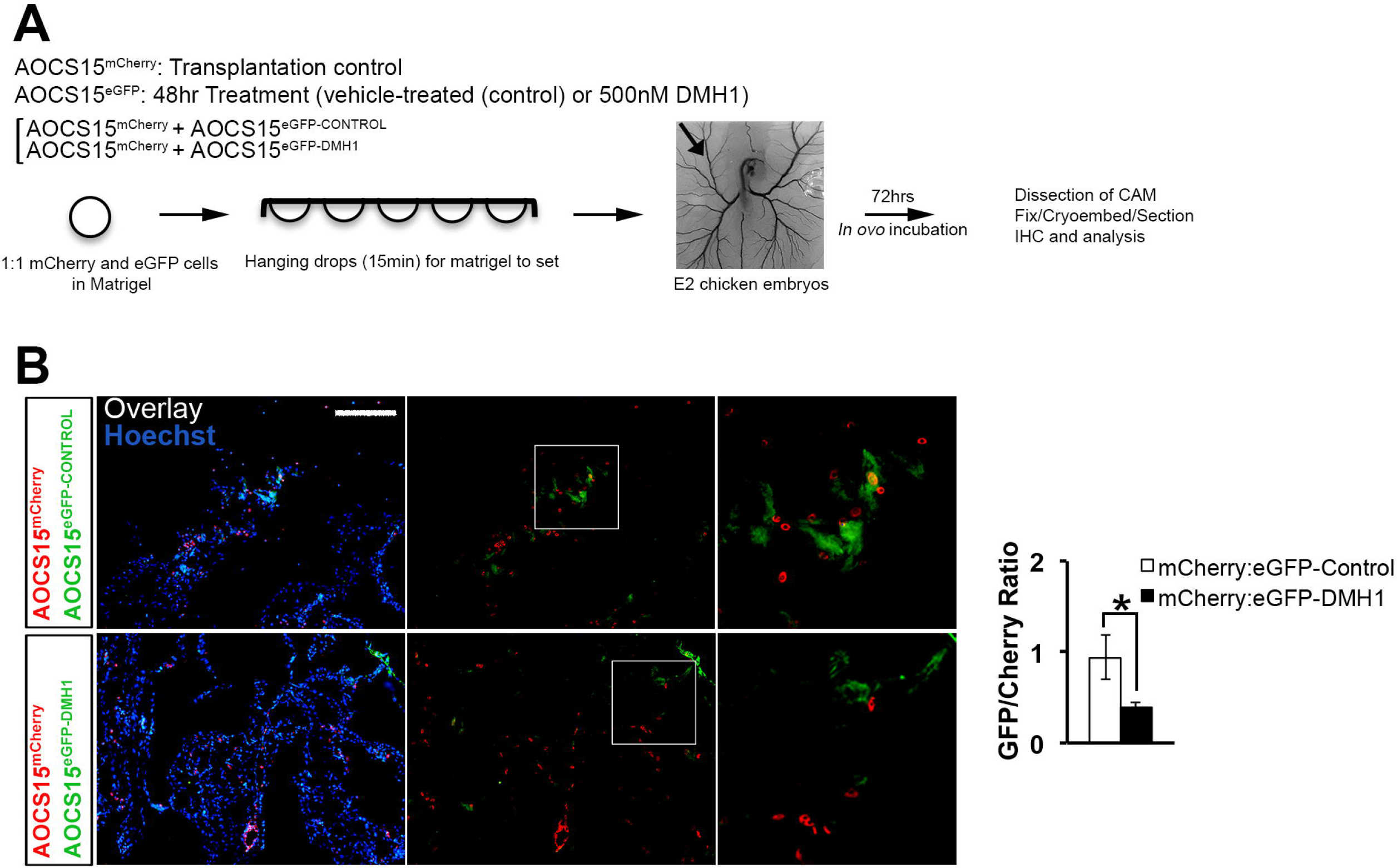
CAM migration assay confirming the role of BMP pathway activity upstream of ovarian tumour cell migration. A) Details of cell transplantation and experimental design. B) Immuno-histochemistry for mCherry and eGFP in CAM samples from control and DMH1- treated cell transplantations (scale bar: 100 μm).

### Ovarian cancer cells maintain a reservoir of autonomous BMP signalling pathway activity

Given the high expression levels of EMT transcription factors in both COLO316 and AOCS15 cells and the confirmed role of BMP pathway activity in the maintenance of the EMT signature, we questioned if the cells maintained a cell-autonomous mode of BMP pathway activity. Both cell lines expressed nuclear pSMAD1/5/9, marking an endogenous level of BMP pathway activity in normal culture conditions (Figure 5A). Further analysis of the cells for the expression of BMP ligands confirmed an endogenous expression for multiple BMP ligands in both COLO316 (*BMP1,4,5,6,8,11,13,14*) and AOCS15 (*BMP1,6,11*) cells (Figure 5B). To confirm the auto/paracrine secretion of the BMP ligands, we blocked BMP6, the ligand commonly expressed between the two cell lines, with a monoclonal antibody and checked for levels of endogenous pSMAD1/5 expression. Treatment with a monoclonal antibody against BMP6 resulted in a significant decrease in the endogenous pSMAD1/5 expression in both cell lines compared to Control and IgG-treated samples (Figure 5C). Together, these findings confirm a potent reservoir of endogenous BMP pathway activity in both cell lines.

**Figure 5.**
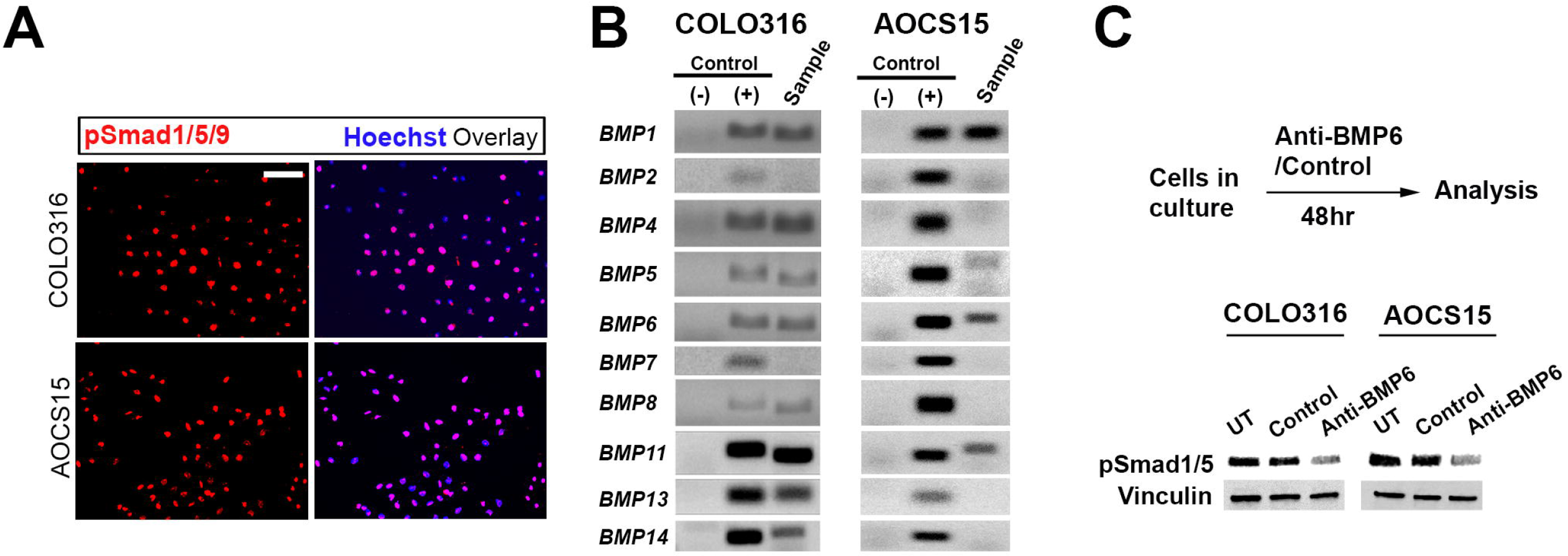
Cell-autonomous BMP pathway activity in ovarian carcinoma cells. A) Immuno- cytochemistry for pSmad1/5/9 in COLO316 and AOCS15 ovarian carcinoma cells (scale bar: 50 μm). B) Semi-quantitative RT-PCR for selected BMP ligands in both cell lines ((+): universal total human RNA, positive control, (-): control for genomic DNA contamination; PCR reaction mix without reverse transcriptase). C) Blocking of secreted BMP6 ligand by a monoclonal antibody for 48 hrs or control IgG followed by Western Blot analysis on total protein for pSmad1/5 expression. Vinculin was used as loading control.

### BMP is the maintenance pathway of choice for EMT in ovarian carcinoma cells secondary to EMT induction

Stratification of signalling pathways along the EMT axis suggested a discrete role for the BMP pathway activity in addition to the previously defined roles as an inducer of EMT [19, 20, 33, 53]. We therefore asked if independent induction of EMT would result in selective cell-autonomous activation of the BMP pathway. For this, two single cell-derived subclones of AOCS15 ovarian carcinoma cells with relatively low levels of pSMAD1/5 expression were selected (Clone#1 and Clone#2; Figure S3E). Treating both clones with the global EMT inducer TNF-α [54, 55] up to 72 hrs, resulted in partial induction of EMT signatures (Figure 6A). Clone#1 showed significant induction of *SNAI1(Snail)* and *TWIST1* (48 and 72 hrs), whereas Clone#2 showed a significant induction of *ZEB1* and *Vimentin* at both 48 and 72 hrs time points. Further analysis for the expression of the downstream BMP pathway activity marker (pSmad1/5) in both clones confirmed elevated levels of pSmad1/5 expression, respectively for Clone#1 and Clone#2, starting from 24 hr and 48 hrs after treatment with TNF-α (Figure 6B). This finding demonstrates a selective, cell-autonomous role for the BMP pathway activity, downstream of EMT induction (Figure 6C).

**Figure 6.**
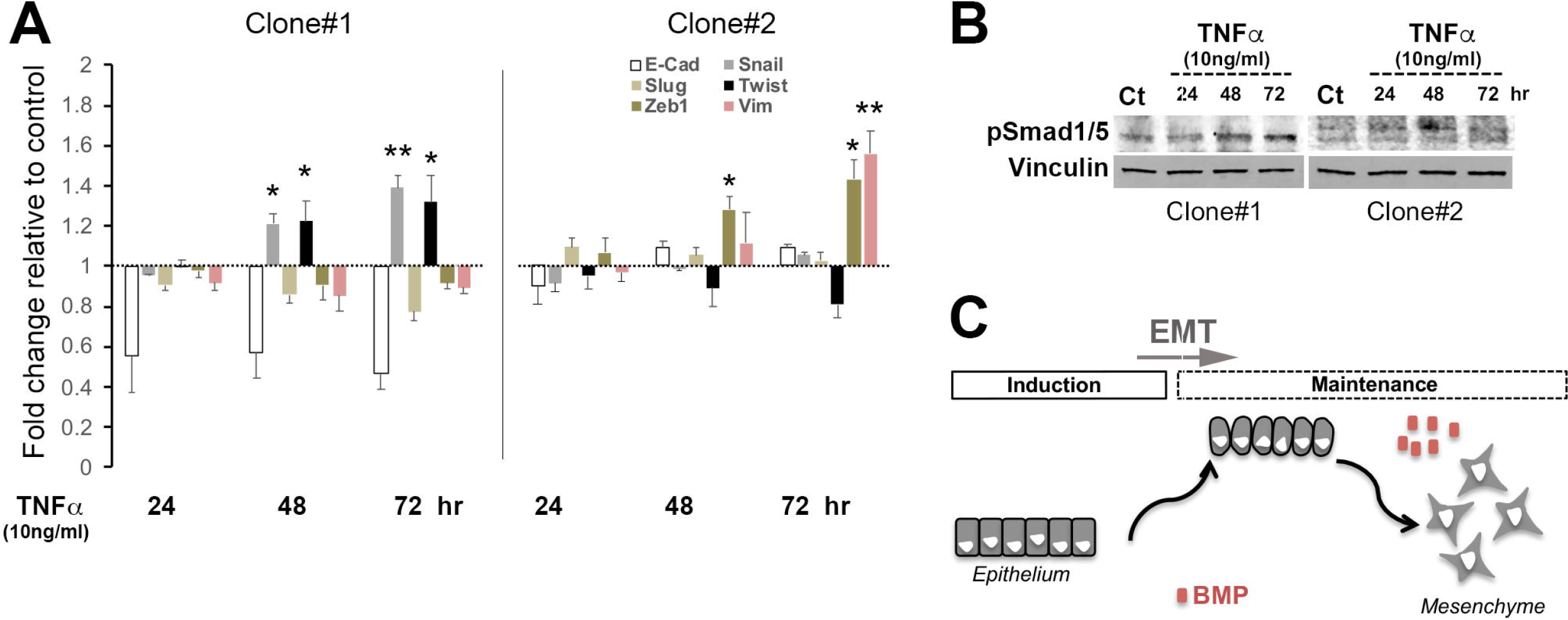
Selective activation of the BMP pathway downstream of EMT induction. A) Real- time RT-PCR on total RNA from two subclones of AOCS15 (Clones #1& #2), for EMT transcriptions factors and E-cadherin (E-CAD), post TNFα treatment. Changes are illustrated relative to the control/vehicle-treated cells. B) Western blot analysis on total protein from each clone for pSMAD1/5. Vinculin was used as loading control. C) Illustration of the proposed mode of action for the BMP pathway during maintenance of the EMT network.

## Discussion

Our study has uncovered a new aspect of the BMP signalling, as a cell-autonomous pathway upstream of EMT maintenance, secondary to EMT induction. Increased expression levels of BMP ligands have been reported in a range of tumours [34, 56–59]. Functionally however, BMP pathway has been defined to have both pro and anti-tumorigenic roles and a range of target programs including primary tumour initiation, EMT and invasion/metastasis [24, 25].

Several studies have defined a role for the BMP pathway in induction of the EMT and tumour cell migration programs. BMP2,4 and 7 induced EMT in human pancreatic cancer cells (Panc- 1) partly through induction of matrix metalloproteinase (MMP)-2 [18]. In ovarian cancer cells, BMP4 induced EMT through activation of Rho GTPase [19]. Further, BMP4-induced EMT was shown to be largely affected by matrix rigidity and the activation of YAP1-CDK8/CDK19 pathway [20]. Inhibition of BMP pathway was as well shown to impair migration and invasion of tumour cells in hepatocellular carcinoma [56]. In breast cancer, BMP2 induced EMT and breast cancer stemness through hyperphosphorylation of the global tumour suppressor protein retinoblastoma (Rb) and induction of CD44 [33, 60]. Likewise, BMP was suggested as a pro-metastatic factor to promote bone remodelling facilitating the metastasis of breast cancer cells into the bone [24]. Despite the positive correlation between BMP pathway activity and pro-invasive cellular programs however, the initial stage of metastasis was shown to require a transient ZEB1-induced generation of BMP inhibitors leading to BMP pathway suppression [61]. Likewise, in cholangiocarcinoma, BMP7 blocked TGFβ-induced cell migration and EMT during the inflammatory response associated with tumourigenesis [62].

The conflicting roles of BMP pathway in tumour cell migration and EMT, on one hand suggests a context-dependant functional outcome for BMP pathway activity with possible additional synergistic/antagonistic pathways. On the other hand, with majority of studies so far, focusing on the role of BMP pathway in the induction of EMT and in light of new insight on EMT as a continuum of semi-stable states [48, 63–66], the question remains if the BMP pathway could be involved in ensuring the stability of any of the intermediate E/M states, or transition from one intermediary state to the other. In our analysis of the TCGA tumour data from high grade serous ovarian cancer, a significant correlation was found between expression levels of EMT markers and that of the two BMP family members, BMP1 and 4, where the former in fact acts as a non-ligand protease with proven functions within the TGF family signalling pathways [67]. The correlating expression trend with EMT transcription factors was as well observed with the target genes of the BMP pathway, the inhibitors of differentiation (ID), and was further stratified as enriched BMP pathway activity in EMT^high^ HGS tumours, suggesting a window of functional activity for this pathway downstream of EMT induction. Further inhibition of endogenous BMP pathway activity by DMH, resulted in a marked reduction of all key EMT-associated transcription factors. However, the repression of EMT program did not result in full diversion from the mesenchymal program or the so-called mesenchymal to epithelial transformation (MET) evidenced by no significant induction of the epithelial marker E-cadherin. It is therefore plausible that additional pathways contribute to the maintenance network upstream of EM, hence impaired BMP pathway alone cannot sufficiently initiate the MET program [68]. Moreover, it is also possible that in the presence of active BMP pathway, the cells have passed through a commitment point (point-of-no-return) and are no longer able to reverse their E/M state. In fact, it is shown that once cells are ‘locked’ in a mesenchymal state by sustained induction of EMT, their reversibility and plasticity can become limited [69, 70]. The sustained, cell-autonomous activity of the BMP signalling pathway, and the cell-voluntary induction of the BMP pathway downstream of TNFα is aligned with the notion that progression along the EMT axis happens in discrete steps fine-tuned by distinct regulatory networks. As such, a state transition is only possible if the existing network acts in synergy with additional cell-autonomous/environmental stimuli to activate the downstream signalling cascade that form the secondary network for the new state. As such, autocrine BMP signals can activate other autocrine and/or paracrine signalling pathways implicated in EMT such as Notch-Jagged signalling [71], which has been implicated in coordinating EMT response at a spatial tissue-level scale [72].

Signalling pathways constitute a crucial component of the regulatory networks upstream of EMT, cancer stemness, metastasis and states/levels of response to therapy and can potentially act to integrate and orchestrate these cellular events. In-depth and detailed knowledge of these pathways will therefore provide fine-tuned targets for development of novel therapeutics and thus potentiate more positive clinical outcomes through targeted therapy.

## Authorship contribution statement

GJ, FZ andTYLL, have contributed to methodology, validation, writing, editing and the review of the original draft. PC and MKJ have contributed to data curation, formal analysis and visualisation. NS, MKJ and AD contributed to conceptualisation, writing, editing and the review of the original draft. JJHC, AOCSG and DDLB have contributed to resources. NS have additionally contributed to the funding acquisition, methodology and project administration.

## Acknowledgements

This study was supported by University of Sydney COMPACT research seed grant, Sydney Dental School research support and the Dr. Poyner award from Australian Dental Research Foundation (ADRF). GJ is funded via a Sydney West Translational Cancer Research Centre (SW-TCRC) PhD Scholarship. SW-TCRC is funded by Cancer Institute NSW. MKJ was supported by Ramanujan Fellowship awarded by Science and Engineering Research Board (SERB), Department of Science and Technology, Government of India (SB/S2/ RJN- 049/2018). JJHC was supported by funding from the National Health and Medical Research Council of Australia (APP1100046). The Australian Ovarian Cancer Study Group was supported by the U.S. Army Medical Research and Materiel Command under DAMD17-01-1-0729, The Cancer Council Victoria, Queensland Cancer Fund, The Cancer Council New South Wales, The Cancer Council South Australia, The Cancer Council Tasmania and The Cancer Foundation of Western Australia (Multi-State Applications 191, 211 and 182) and the National Health and Medical Research Council of Australia (NHMRC; ID199600; ID400413 and ID400281). The Australian Ovarian Cancer Study gratefully acknowledges additional support from Ovarian Cancer Australia and the Peter MacCallum Foundation. The AOCS also acknowledges the cooperation of the participating institutions in Australia and acknowledges the contribution of the study nurses, research assistants and all clinical and scientific collaborators to the study. The complete AOCS Study Group can be found at www.aocstudy.org. We would like to thank all of the women who participated in these research programs.

## Abbreviations

EMT: Epithelial Mesenchymal Transition
BMP: Bone morphogenic protein
SMAD: Small mothers against decapentaplegics
TGF-β: Transforming growth factor-β
PI3K: Phosphatidylinositol 3-kinase
MAPK: Mitogen-activated protein kinase
NFκB: Nuclear factor kappa B
JAK/STAT: Janus kinase/signal transducer and activator of transcription
CSC: Cancer Stem Cell
HGS: High-grade serous
TCGA: The Cancer Genome Atlas
ID: Inhibitor of differentiation
CAM: Chorioalantoic membrane
CFA: Cell-free area
MET: Mesenchymal to epithelial transition

## Supplementary Figures and Tables

**Figure S1.**
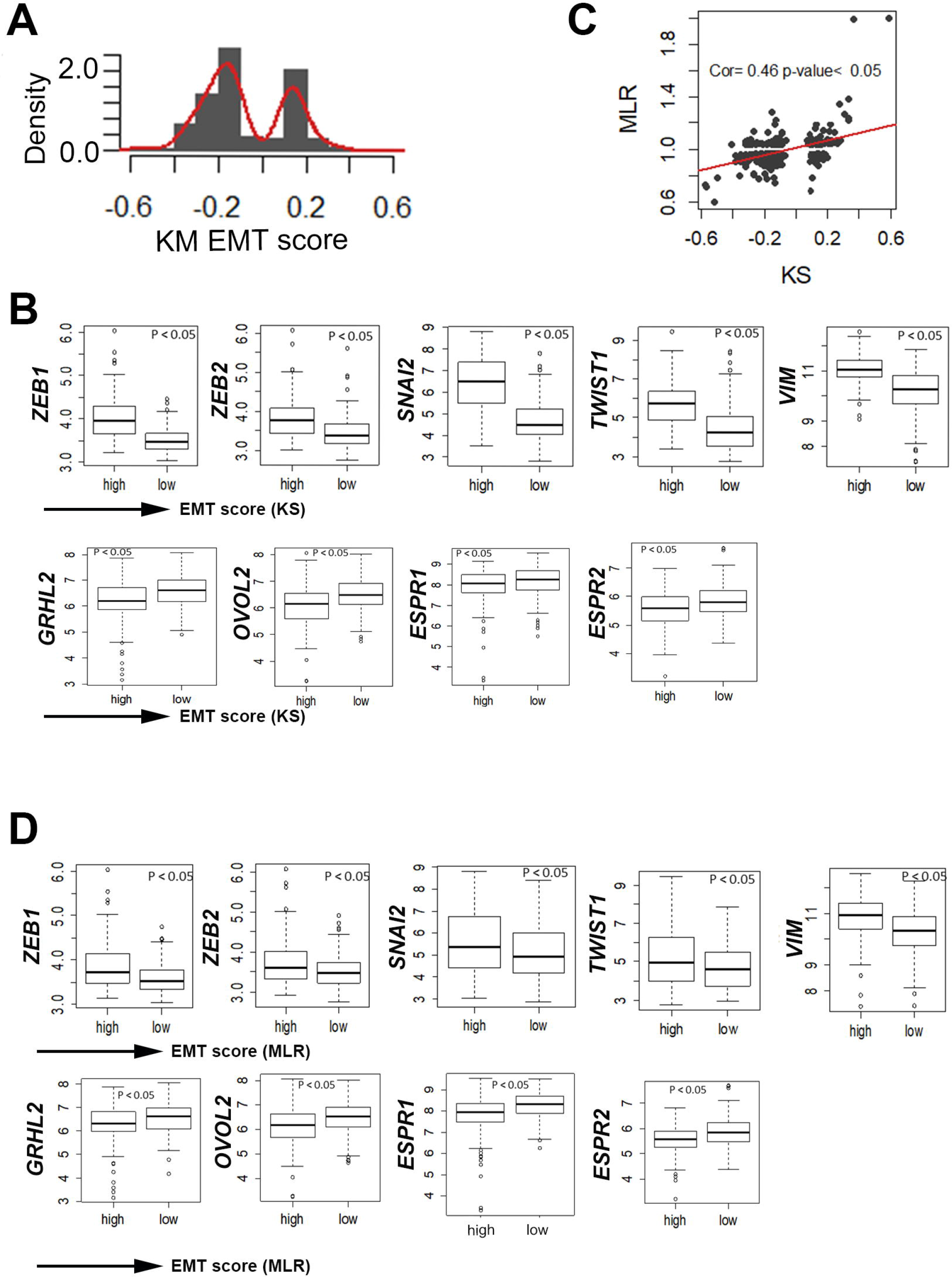
Verification of the EMT scoring methods in HGS ovarian tumour samples. A) Distribution profile of EMT scores based on the KM method among HGSC samples. B) Expression of epithelial or mesenchymal markers in subsets of HGSC tumour samples scored for low and high levels of EMT using the KS method. C) Correlation between EMT scores calculated by two separate methods of KM and MLR. D) Expression of epithelial or mesenchymal markers in subsets of HGS tumour samples scored for low and high levels of EMT using the MLR method.

**Figure S2.**
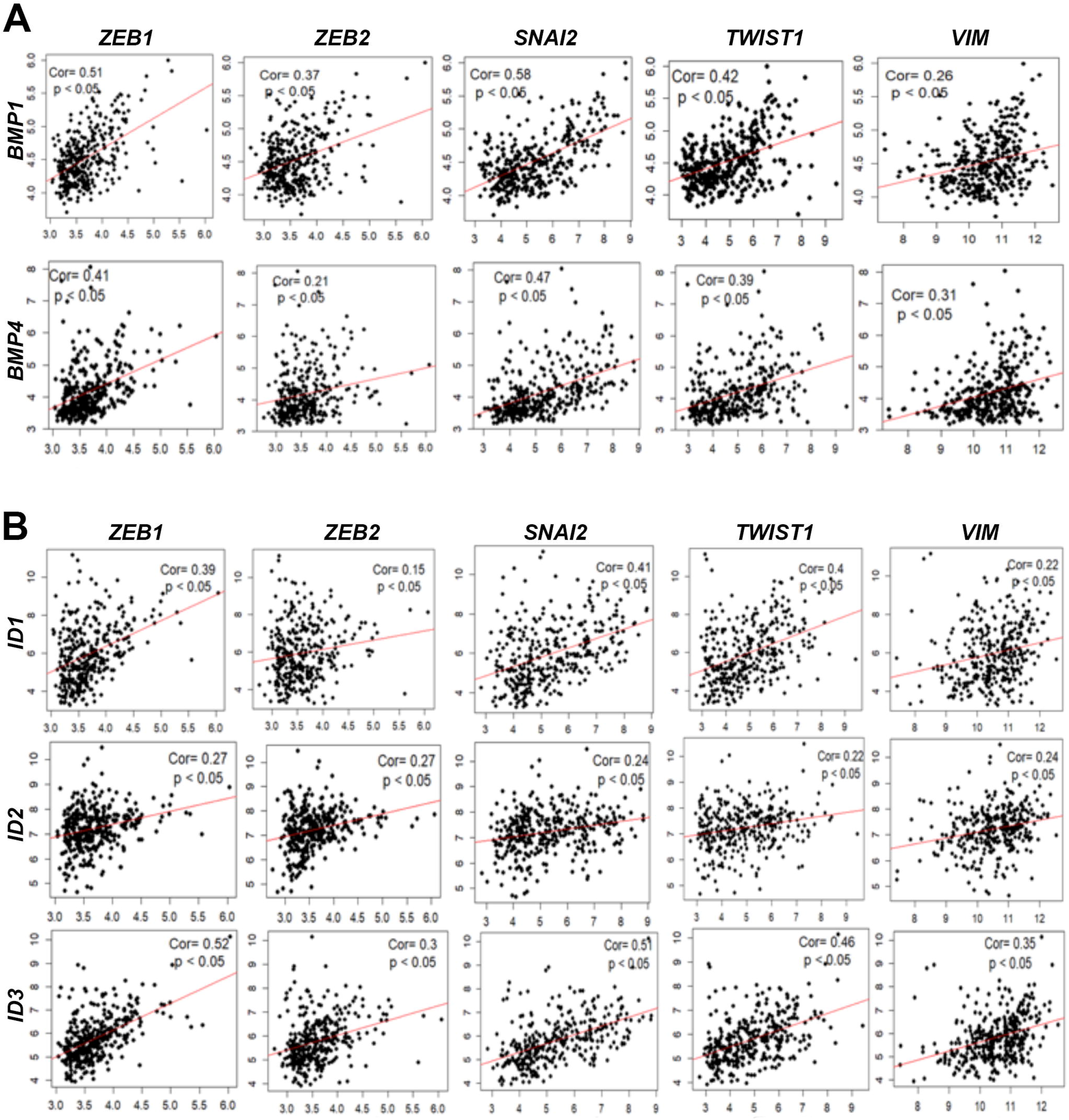
Correlation analysis between the expression of BMP and EMT transcription factors. A) Expression correlation analysis between BMP1 & 4 and EMT transcription factors and the mesenchymal marker Vimentin (VIM). B) Expression correlation analysis between ID1-3 and EMT transcription factors and the mesenchymal marker Vimentin (VIM). The linear regression line is depicted in red.

**Figure S3.**
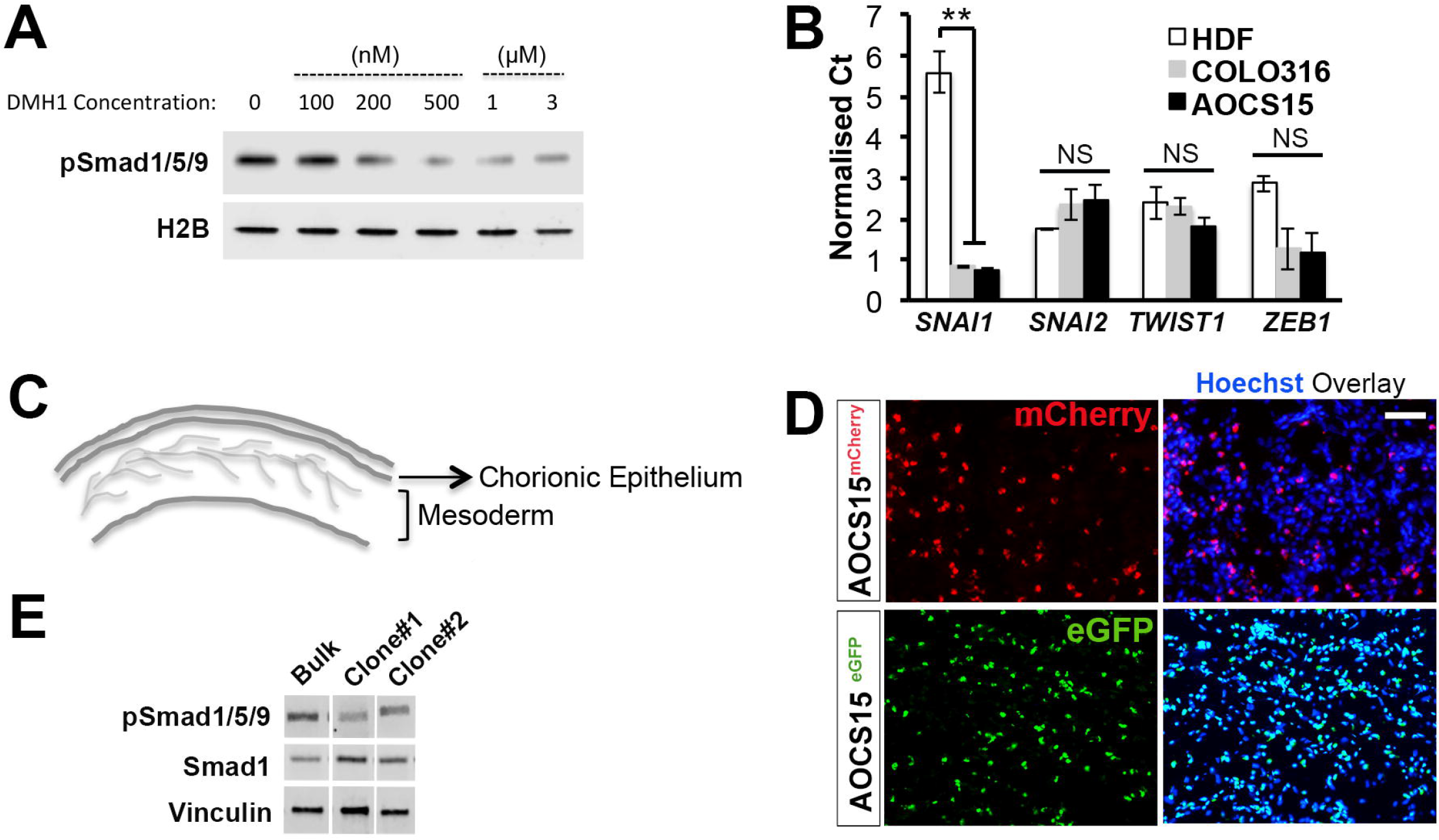
Supporting data for BMP pathway inhibition experiments. A) Western blot analysis on total protein from ovarian cancer cells treated with varying concentrations of DMH1, for the expression of pSmad1/5/9. Histone H2B was used as loading control. B) Real-time RT-PCR for EMT transcriptions factors on total RNA from COLO316 and AOCS15 cells compared to human dermal fibroblasts (HDF) as a control mesenchymal cell (NS: not significant). C) Illustration highlighting the structure of the chicken chorioalantoic membrane (CAM). D) Immune-histochemistry for mCherry and eGFP, in CAM samples transplanted with AOCS^mCherry^ and AOCS^eGFP^ cells (Scale bar: 50 μm). E) Western blot analysis on total protein from AOCS15 bulk cells vs selected subclones (#1 and 2), for the expression of pSMAD1/5/8 and Smad1. Vinculin was used as loading control.

**Table S1:**
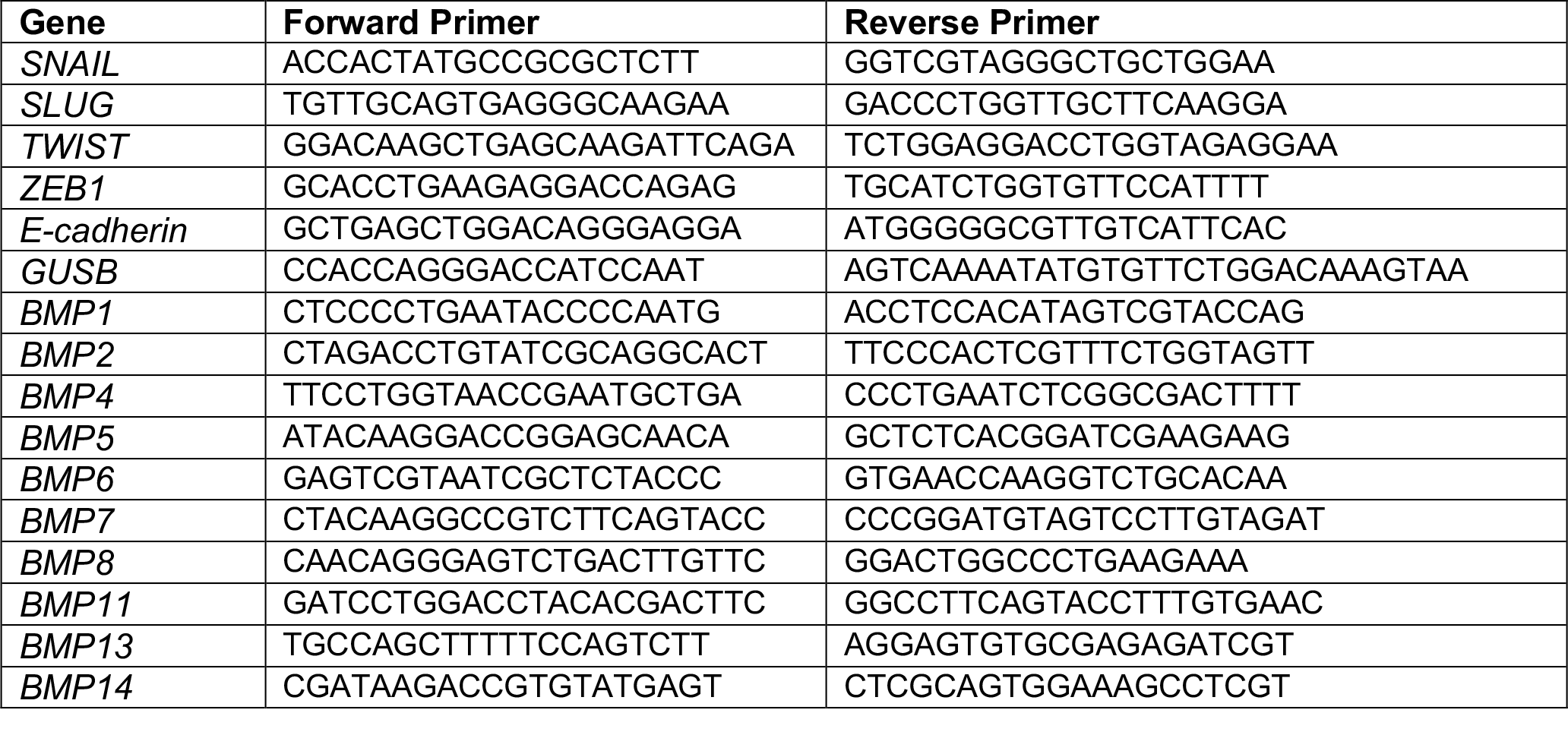
List of gene-specific primers used in Real-time and semi-quantitative RT-PCR.

**Table S2:** Correlation values for epithelial (Epi) and mesenchymal (Mes) markers in tumours with low and high EMT scores.

| **Epithelial**<br>**marker** | **Mesenchymal**<br>**marker** | **Cor** | **p-value** |
| --- | --- | --- | --- |
| OVOL2 | TWIST1 | -0.26 | <0.05 |
| GRHL2 | ZEB1 | -0.24 | <0.05 |
| ESRP1 | ZEB1 | -0.22 | <0.05 |
| CDH1 | TWIST1 | -0.2 | <0.05 |
| ESRP2 | VIM | -0.2 | <0.05 |
| OVOL2 | VIM | -0.18 | <0.05 |
| CDH1 | ZEB1 | -0.17 | <0.05 |
| ESRP2 | ZEB1 | -0.17 | <0.05 |
| CDH1 | ZEB2 | -0.17 | <0.05 |
| ESRP2 | TWIST1 | -0.16 | <0.05 |
| OVOL2 | CDH2 | -0.15 | <0.05 |
| GRHL2 | TWIST1 | -0.13 | <0.05 |
| CDH1 | CDH2 | 0.12 | <0.05 |
| OVOL2 | ZEB2 | -0.11 | <0.05 |

